# Vorinostat corrects cognitive and non-cognitive symptoms in a mouse model of fragile X syndrome

**DOI:** 10.1101/2020.08.05.238568

**Authors:** Qi Ding, Xueting Wu, Xuan Li, Hongbing Wang

## Abstract

Fragile X syndrome (FXS) is caused by mutations in the *FMR1* (fragile X mental retardation 1) gene. It is a significant form of heritable intellectual disability with comorbidity of other symptoms such as autism. Due to the lack of efficacious medication, repurposing the existing FDA-approved drugs may offer an opportunity to advance clinical intervention for FXS. Analysis of the whole-genome transcription signatures predicts new therapeutic action of vorinostat to correct pathological alterations associated with FXS. We further find that the administration of vorinostat restores object location memory and passive avoidance memory in the *Fmr1* knockout (KO) mice. For the non-cognitive behavioral symptoms, vorinostat corrects the autism-associated alterations, including repetitive behavior and social interaction deficits. In the open field test, vorinostat dampens hyperactivity in the center area of the arena. Surprisingly, vorinostat does not affect the abnormally elevated protein synthesis in *Fmr1* KO neurons, suggesting different outcomes from correcting behavioral symptoms and specific aspects of cellular pathology. Our data reveal the therapeutic effects of the FDA-approved drug vorinostat in a mouse model of FXS and advocate efficacy testing with human patients.

## INTRODUCTION

Fragile X syndrome (FXS) is a genetic disorder caused by mutations in the *FMR1* (fragile X mental retardation 1) gene. As the most frequent mutation, the increased number of the CGG trinucleotide repeat in the 5’ non-coding region hampers gene transcription and leads to a significant reduction or lack of FMRP (fragile X mental retardation protein) expression. The main symptoms of FXS patients include cognitive disability, hyperactivity, and autism-related behavior ^1,2^. Based on the pathological and mechanistic studies with animal models of FXS, strategies targeting specific molecular and cellular alterations have shown notable therapeutic efficacy in preclinical studies ^3-7^. Nevertheless, efficacious medication is not available yet.

Toward achieving clinical treatment for FXS, the development of brand-new drugs may easily take more than ten years. Comparing to *de novo* drug development, repurposing the existing FDA-approved drugs, of which the toxicity has already been tested in humans, and the detailed pharmacology and formulation are available, provides unprecedented opportunity to cut the amount of money, time, and effort. The main traditional approaches of drug repurposing usually depend on the outcome of high-throughput screening ^8^ or knowledge of drug structure and mechanism of action. More recently, computational comparison of drug-induced transcriptome profiles has been proposed as a non-structure based *in-silico* screening of similarity drugs ^9-11^. By using such a computation approach, our recent study predicts that the FDA-approved drug vorinostat may have therapeutic effects to correct FXS-associated symptoms ^12^. However, the prediction requires empirical validation.

Vorinostat, also known as suberanilohydroxamic acid (SAHA), is currently used to treat cancers, including cutaneous T cell lymphoma (CTCL) and advanced non-small-cell lung carcinoma (NSCLC). Although, as a small chemical compound, it should have promiscuous pharmacological activities, its inhibition activity against class I, II, and IV histone deacetylases (HDAC) is well recognized. Vorinostat also shows neuroprotective effects in the central nervous system and is suggested to treat neurodegeneration such as Alzheimer’s disease and Huntington disease ^13-15^. As histone acetylation-induced epigenetic changes are involved in activity-dependent plasticity and learning ^16^, a well-recognized outcome of vorinostat treatment is the enhancement of memory and cognition ^15^.

In this study, we examined the effects of vorinostat in the *Fmr1* KO mice. We found that vorinostat corrected deficits in object location memory and passive avoidance memory. Interestingly, it also corrected non-cognitive behavioral symptoms, including repetitive behavior, social interaction deficits, and a specific aspect of hyperactivity. Our study provides evidence to support a new therapeutic action of vorinostat that is predicted by an unbiased transcriptome-based computational approach. It also advocates future tests with human clinical trials.

## MATERIALS AND METHODS

### Comparison of vorinostat-induced transcriptome changes with other drugs in the Connectivity Map (CMap) database

The search for drugs/compounds that induce similar transcriptome changes to that of vorinostat in the CMap database was performed as described ^12^. In brief, we first obtained microarray data sets of MCF7, PC3, and HL60 cells treated with vorinostat and the corresponding vehicle controls. The differential gene expression analysis was conducted, and the top 500 up-regulated and down-regulated genes were retained as the signature for further query ^12,17,18^. The transcriptome signature of vorinostat was uploaded to the CMap query page and used to search for compounds that induce similar or oppositional transcriptome changes in the corresponding cell line (i.e., MCF7, PC3, and HL60) ^9^. The returning results of drugs/compounds with a p-value of less than 0.05 in each cell line are presented in Supplementary Tables 1, 2, and 3.

### Animals

Young adult male mice at 2.5 to 3.5 months of age were used for behavioral examinations. Postnatal day 0 mice were used to obtain primary hippocampal neurons. The *Fmr1* knockout (KO) and their wild type (WT) littermates are on the C57BL/6 background. The Institutional Animal Care and Use Committee approved all procedures.

### Behavioral examinations

For the examination of object location memory, mice were habituated to the training chamber without any object for 10 min on three consecutive days. Mice were then trained by a 10-min exposure to the training chamber holding two objects at different locations (Fig. 2a), during which the mice freely explored the chamber and interacted with the objects. Twenty-four hours after training, the trained mice were tested by a 10-min re-exposure to the same chamber with one object at the same location and one object at a new location (Fig. 2a). During training and testing, time spending in interacting with each object was recorded. The % of preference was determined by the time spent with each object divided by the total time spent with both objects.

**Figure 1.**
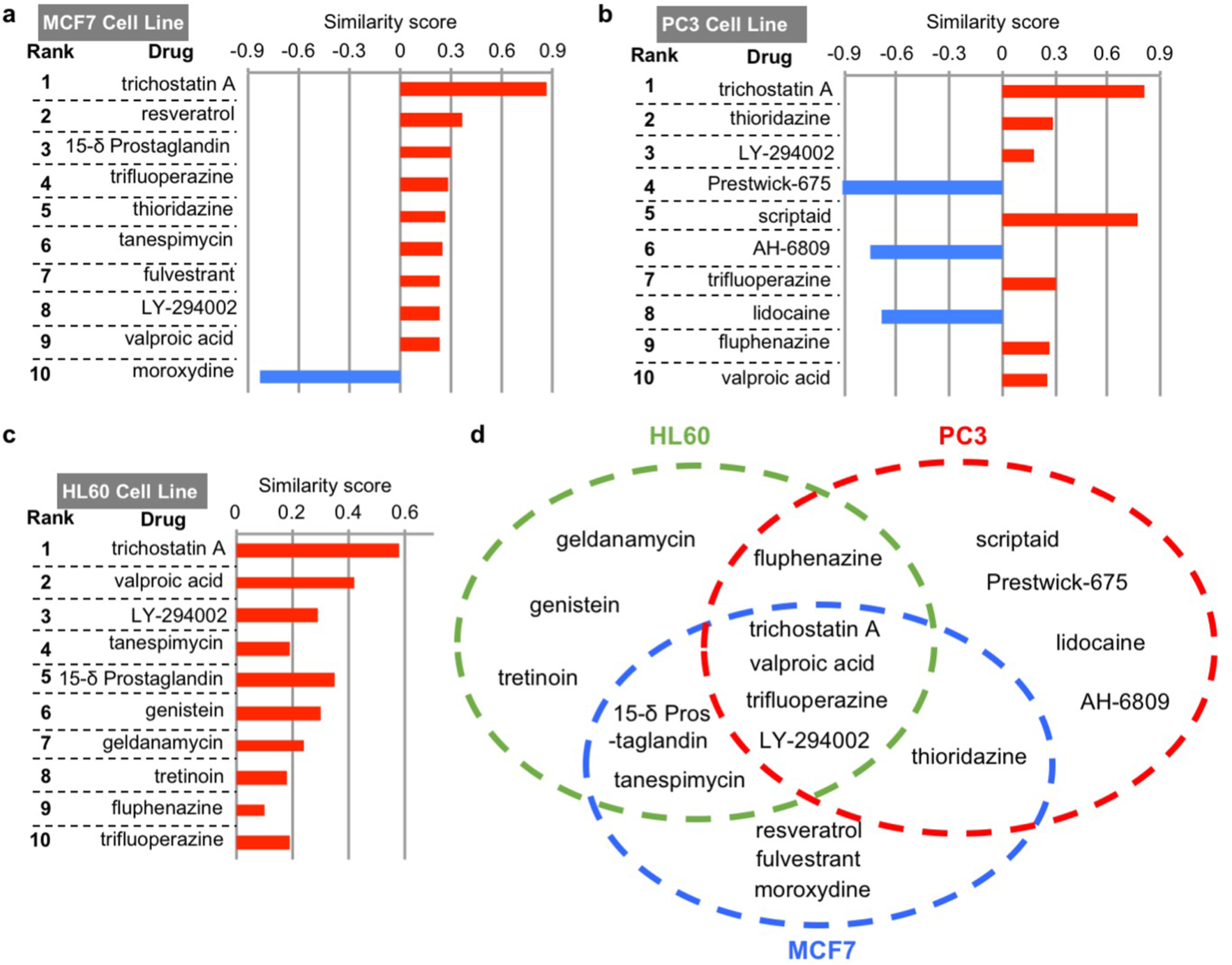
Similarity drugs of vorinostat identified by the drug-induced transcriptome changes. The top 10 similarity drugs/compounds of vorinostat, along with the similarity scores, in MCF7, PC3, and HL60 cell lines, are shown in **a, b**, and **c**, respectively. Some similarity drugs/compounds of vorinostat are identified from a unique cell line, as indicated in **d**. Some similarity drugs/compounds induce similar transcriptome changes to that of vorinostat in multiple cell lines (**d**).

**Figure 2.**
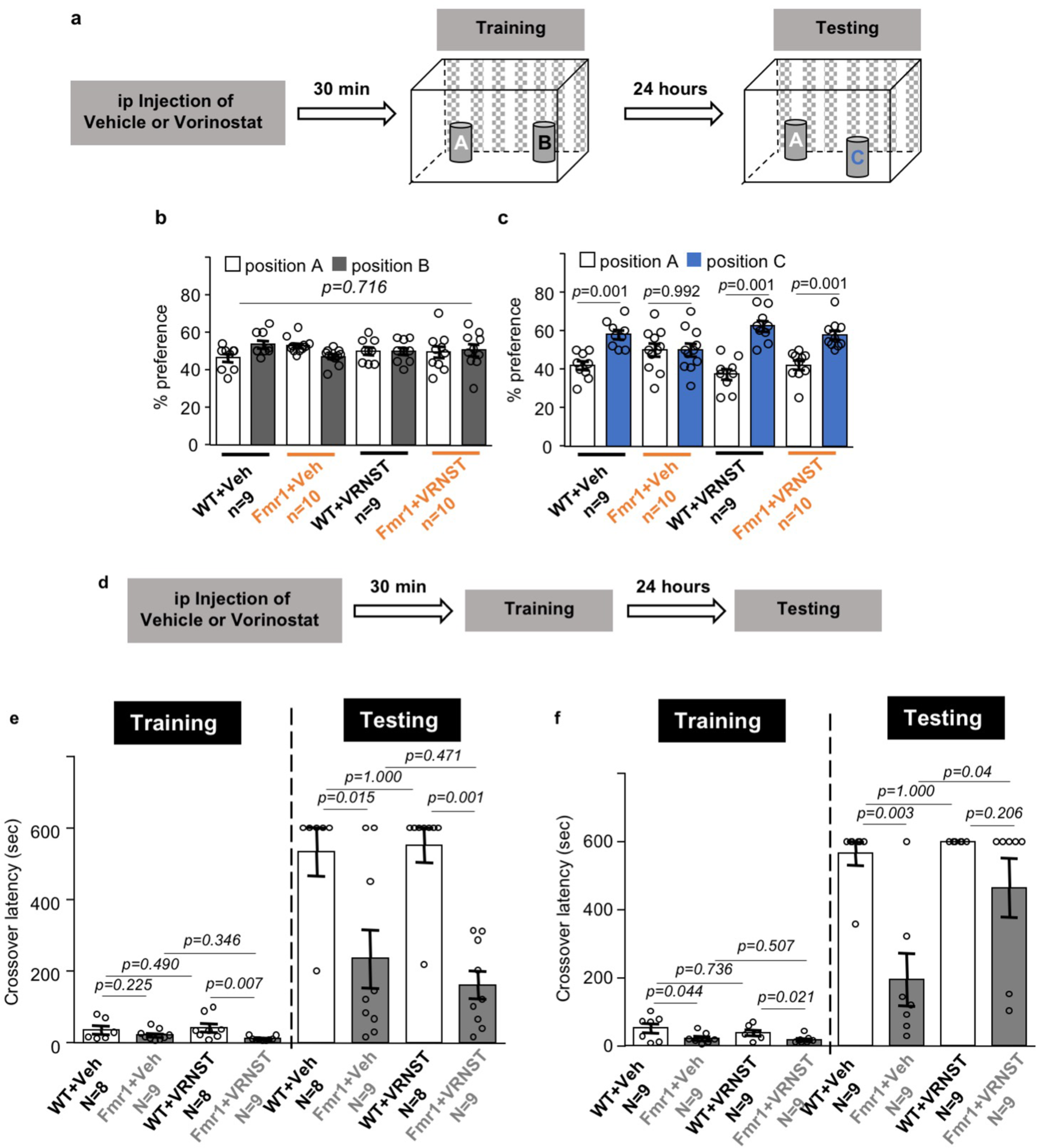
Effects of vorinostat on the correction of object location memory and passive avoidance memory in the *Fmr1* KO mice. **a**. Drug administration and object location memory paradigm. Following a single ip injection with vehicle (Veh) or vorinostat (VRNST), wild type (WT) and *Fmr1* KO (Fmr1) mice were subject to a training chamber with certain spatial cues and allowed to explore two objects at location A and B. During testing, the trained mice were reintroduced to the training chamber and allowed to interact two objects at location A and C. **b**. Mouse preference to the object at locations A and B during training. **c**. Mouse preference to the object at locations A and C during testing. **d**. Timeline of drug administration, training, and testing for passive avoidance memory. Wild type (WT) and *Fmr1* KO (Fmr1) mice were first injected with vehicle (Veh) or vorinostat (VRNST) and then received passive avoidance training. The trained mice were tested 24 hours later. **e**. Mice were trained after a single injection of Veh or VRNST. **f**. Mice were first injected with Veh or VRNST daily for two weeks and then trained after the last injection. Crossover latency during training and testing was recorded (**e** and **f**). Data are presented as mean +/- SEM. The *p*-values in **b** and **c** were determined by three-way ANOVA followed by *post hoc* pairwise comparison. The *p*-values for passive avoidance training data (**e** and **f**) were determined by two-way ANOVA followed by *post hoc* pairwise comparison. The *p*-values for the passive avoidance testing data (**e** and **f**) were determined by the Fisher exact test.

For the examination of passive avoidance memory, mice were trained as described in our previous studies ^19^. During training, mice received a mild foot shock (0.7 mA for 2 sec) immediately after entering the dark chamber, stayed in the dark chamber for 30 sec, and were then returned to their home cage. Twenty-four hours later, the trained mice were reintroduced to the training chamber; crossover latency (i.e., the time elapsed until the mice crossed over and entered the dark chamber) was recorded. When there was no crossover beyond 600 sec, the examination was terminated, and a crossover latency of 600 sec was used for those specific mice.

Mouse activity in the light/dark box, the open field, and the 3-chamber social interaction test was examined as described in our previous studies ^3,12,19^.

### Drug administration

Vorinostat (Sigma-Aldrich) and trichostatin A (Sigma-Aldrich) were prepared in the vehicle (10% DMSO) and i.p. injected into mice at 50 mg/kg and 10 mg/kg, respectively. The chosen doses were equivalent or higher than doses used in previous *in vivo* studies, showing that such treatments are sufficient to improve cognitive functions in mice ^16,20-22^. For the examination of object location and passive avoidance memory, drugs were administered 30 min before training. For all other behavioral paradigms, drugs were administered 30 min before the examination. The control groups were treated with vehicle injection.

### Examination of protein synthesis and histone acetylation in neurons

Primary hippocampal neurons were obtained from postnatal day 0 WT and *Fmr1* KO mice and maintained *in vitro*. To determine protein synthesis with the SUnSET method ^23^, DIV (days *in vitro*) 14 neurons were incubated with 5µg/ml puromycin (Sigma, Cat #P8833) for 30 min and then harvested in Buffer H (50 mM β-glycerophosphate, 1.5 mM EGTA, 0.1 mM Na_3_VO_4_, 1 mM DTT). After the determination of total protein concentration, the samples were examined by Western blot with anti-puromycin antibody (KeraFAST, Cat # EQ0001, 1:1000). For the determination of histone acetylation level, samples collected from DIV 14 WT and *Fmr1* KO neurons were examined by Western blot with antibodies against total and acetylated histone proteins H2B, H3, and H4 (Cell Signaling, 1:1000). The relative amount of loading was determined by β-actin. The intensity of the immuno-signal was analyzed by the ImageJ software (NIH, MD, USA).

### Data collection and statistics

Mice were randomly assigned to vehicle and drug treatment groups, which were not disclosed before data analysis. Mice from multiple litters were used to avoid pseudo repeats. Data with normal distribution were analyzed by two-sided Student’s *t*-test or ANOVA. The crossover latency data for passive avoidance testing did not show normal distribution and were analyzed by Fisher’s exact test. All data are expressed as mean ± SEM.

## RESULTS

### *In-silico* screening of vorinostat similarity drugs in the CMap database

Transcriptome landscape reflects a particular aspect of molecular outcome in physiological and pathological conditions ^24-26^. We recently found that transcriptome changes in the *Fmr1* KO neurons can successfully predict therapeutic interventions. Among the predicted drugs, an FDA-approved antipsychotics trifluoperazine causes transcriptome alteration oppositional to that caused by FMRP deficiency. It was further demonstrated that trifluoperazine corrects the key FXS-associated symptoms in the *Fmr1* KO mice ^12^. Moreover, computational analysis of the trifluoperazine-induced transcriptome signature revealed other similarity drugs, predicting that those similarity drugs may be repurposed to treat FXS. Among the top 10 similarity drugs, vorinostat is the third-ranked FDA-approved drug following two antipsychotics ^12^.

Here, we further used vorinostat-induced transcriptome perturbations as a query to identify similarity drugs in the CMap database (Broad Build 02 database, http://www.broadinstitute.org), which contains over 7000 reference gene signatures altered by 1309 compounds/perturbagens. As the gene signatures were characterized in three major cell lines (i.e., MCF7, PC3, and HL60) in the database, we performed a computational analysis to search for similarity drugs within each cell line. The transcriptome signature of vorinostat in each cell line identified drugs showing significant positive and negative similarity scores (i.e., similarity mean with a *p*-value of less than 0.05) (Supplementary Table 1, 2, and 3). Notably, there is an overlap of similarity drugs among the three cell lines, indicating a certain degree of conservation of transcriptional responses to vorinostat. Among the top 10 ranked compounds (Fig. 1a, 1b, and 1c), 4 drugs are the common vorinostat similarity drugs identified from all 3 cell lines (Fig. 1d). These 4 similarity drugs include 2 HDAC inhibitors (i.e., trichostatin A and valproic acid), trifluoperazine, and a known PI3K (phosphatidylinositol 3-kinase) inhibitor LY-294002. These data further support that, based on their effects on transcriptome signature, vorinostat and trifluoperazine may have a similar action. To validate the *in-silico* prediction of drug action, we examined the effects of vorinostat in a mouse model of FXS.

### Vorinostat corrects cognitive deficits in the *Fmr1* KO mice

Recapitulating the intellectual deficits in human FXS patients, the *Fmr1* KO mice show compromised cognitive function ^2,19^. We first examined the effects of vorinostat on object location memory (Fig. 2a). During training, the vehicle- and vorinostat-treated WT and *Fmr1* KO mice showed comparable preference to objects at location A and B (genotype effect: *F*_1,70_ = 0.001, *p* = 1.000; drug effect: *F*_1,70_ = 0.001, *p* = 1.000; location effect: *F*_1,70_ = 0.133, *p* = 0.716; Fig. 2b). During testing, there was a significant effect of location preference (genotype effect: *F*_1,70_ = 0.001, *p* = 1.000; drug effect: *F*_1,70_ = 0.001, *p* = 1.000; location effect: *F*_1,70_ = 53.032, *p* = 0.001; Fig. 2c). The vehicle-treated WT but not *Fmr1* KO mice showed preference to the object at the new location (i.e., location C), indicating that object location memory is compromised in *Fmr1* KO mice (Fig. 2c). In contrast, both the vorinostat-treated WT and *Fmr1* KO mice showed preference to the object at location C (Fig. 2c), indicating significant location memory formation.

We next examined the effects of vorinostat on passive avoidance memory (Fig. 2d). During training, there was a genotype effect but no drug effect on crossover latency (genotype effect: *F*_1,30_ = 8.514, *p* = 0.007; drug effect: *F*_1,30_ = 0.022, *p* = 0.884; genotype X drug interaction: *F*_1,30_ = 1.358, *p* = 0.253; Fig. 2e). The vehicle-treated wild type (WT) and *Fmr1* KO mice showed similar crossover latency during training (Fig. 2e). Following a single vorinostat administration, *Fmr1* KO mice showed less crossover latency than WT mice (Fig. 2e) during training. Still, there is no significant difference between vehicle- and vorinostat-treated *Fmr1* KO mice (Fig. 2e). When tested 24 hours later, the vehicle-treated *Fmr1* KO mice showed less crossover latency than the vehicle-treated WT mice (Fig. 2e), indicating impaired passive avoidance memory. The acute and single administration of vorinostat did not improve memory in the *Fmr1* KO mice (Fig. 2e). We next treated mice with repeated vorinostat administration by daily intraperitoneal (i.p.) injection for 2 weeks. There was a genotype effect but no drug effect on behavior during training (genotype effect: *F*_1,32_ = 10.201, *p* = 0.003; drug effect: *F*_1,32_ = 0.511, *p* = 0.480; genotype X drug interaction: *F*_1,32_ = 0.054, *p* = 0.817; Fig. 2f). Repeated vorinostat restored passive avoidance memory in the *Fmr1* KO mice to the WT level, as indicated by the improved crossover latency during testing (Fig. 2f).

### Therapeutic effects of vorinostat on repetitive behavior and hyperactivity in the *Fmr1* KO mice

In the light/dark box test, the vehicle-treated *Fmr1* KO mice showed a higher number of repetitive transitions between the light and dark chambers (Fig. 3a1). Administration of vorinostat normalized this hyperactive and repetitive behavior in the *Fmr1* KO mice (genotype effect: *F*_1,39_ = 12.932, *p* = 0.001; drug effect: *F*_1,39_ = 4.469, *p* = 0.041; genotype X drug interaction: *F*_1,39_ = 4.364, *p* = 0.043; Fig. 3a1). Regardless of the treatment, the *Fmr1* KO and WT mice showed similar preference and spent comparable time in the light chamber (genotype effect: *F*_1,39_ = 0.156, *p* = 0.695; drug effect: *F*_1,39_ = 0.042, *p* = 0.840; genotype X drug interaction: *F*_1,39_ = 5.410, *p* = 0.025; Fig. 3a2).

**Figure 3.**
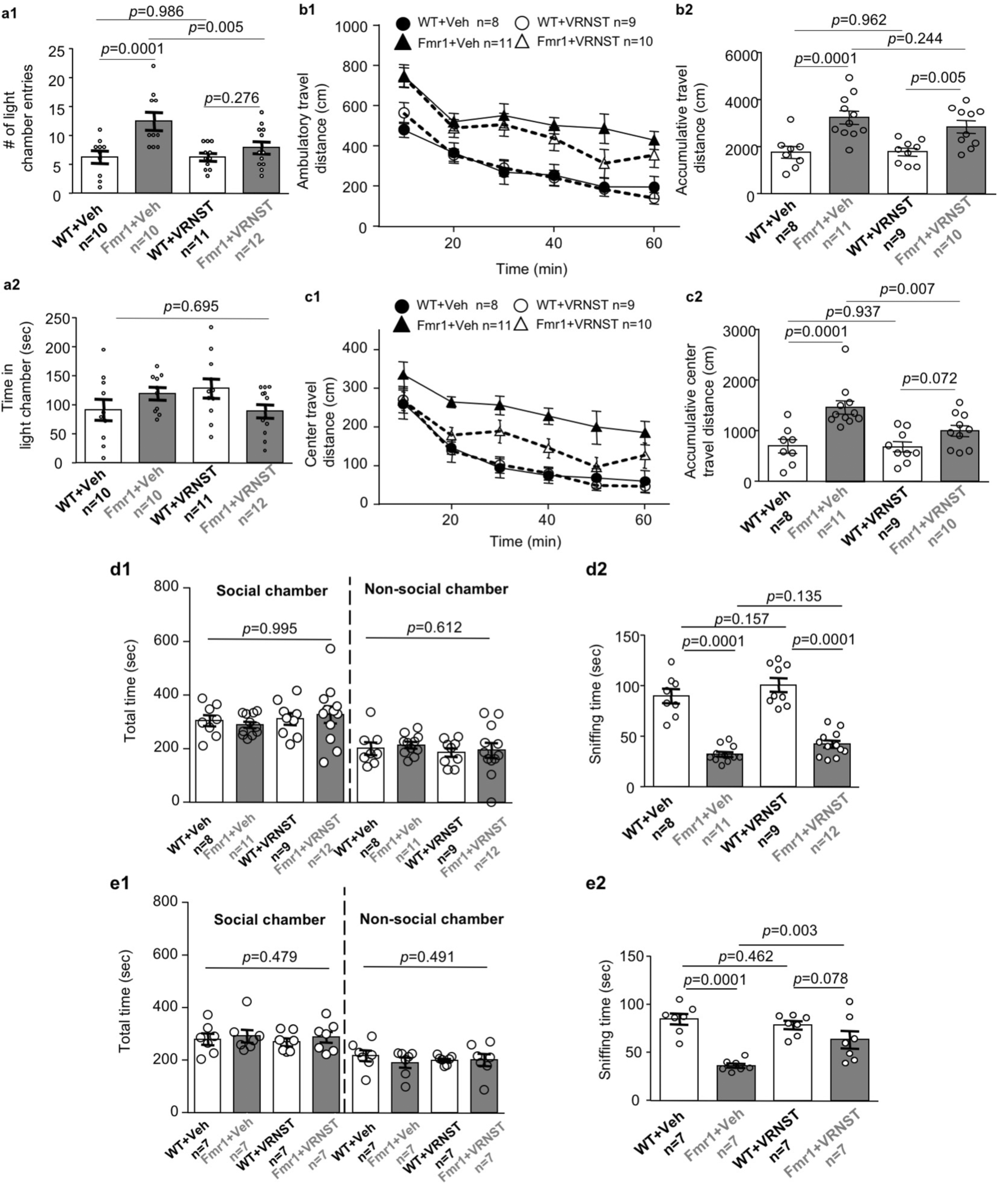
Effects of vorinostat on repetitive behavior, hyper-locomotion, and social interaction deficits in the *Fmr1* KO mice. Following a single (**a, b, c**, and **d**) or repeated (daily for two weeks) (**e**) administration with vehicle (Veh) or vorinostat (VRNST), wild type (WT) and *Fmr1* KO (Fmr1) mice were subject to the light/dark box test (**a**), the open field test (**b** and **c**), and the 3-chamber social interaction test (**d** and **e**). **a**. During the light/dark box test, the mice were allowed to make transitional moves between the light and the dark chamber. The number of entries to the light chamber (**a1**) and time spent in the light chamber (**a2**) are presented as mean +/- SEM. **b** and **c**. During the 60-min open field test, ambulatory travel distance within the whole arena (**b**) and in the center area (**c**) was recorded. Locomotion for each of the 10 min bin (**b1** and **c1**) and accumulative locomotion during the whole 60 min testing (**b2** and **c2**) are presented as mean +/- SEM. **d** and **e**. During the 10-min social interaction test, the total time spent in the social and non-social chamber was recorded, and data are presented in **d1** and **e1**. Time spent in direct interaction with the stranger mouse is shown in **d2** and **e2**. Data are presented as mean +/- SEM. The *p*-values were determined by two-way ANOVA followed by *post hoc* pairwise comparison.

In the open field test, the vehicle-treated *Fmr1* KO mice showed more locomotion activity than the vehicle-treated WT mice in the whole arena (genotype effect: *F*_1,34_ = 25.844, *p* = 0.001; drug effect: *F*_1,34_ = 0.573, *p* = 0.454; genotype X drug interaction: *F*_1,34_ = 0.685, *p* = 0.414; Fig. 3b). They also showed higher locomotion activity in the center area (genotype effect: *F*_1,34_ = 19.996, *p* = 0.001; drug effect: *F*_1,34_ = 3.910, *p* = 0.056; genotype X drug interaction: *F*_1,34_ = 3.457, *p* = 0.072; Fig. 3c). Following vorinostat administration, the *Fmr1* KO mice did not show changes of locomotion in the whole arena (Fig. 3b); they showed a reduction of locomotion in the center area (Fig. 3c).

### Therapeutic effects of vorinostat on social deficits in the *Fmr1* KO mice

In the 3-chamber social interaction test, all groups of mice spent similar time in the social chamber (genotype effect: *F*_1,36_ = 0.000, *p* = 0.995; drug effect: *F*_1,36_ = 0.858, *p* = 0.360; genotype X drug interaction: *F*_1,36_ = 0.443, *p* = 0.510; Fig. 3d1)(genotype effect: *F*_1,24_ = 0.518, *p* = 0.479; drug effect: *F*_1,24_ = 0.122, *p* = 0.730; genotype X drug interaction: *F*_1,24_ = 0.033, *p* = 0.857; Fig. 3e1) as well as in the non-social chamber (genotype effect: *F*_1,36_ = 0.262 *p* = 0.612; drug effect: *F*_1,36_ = 0.695, *p* = 0.410; genotype X drug interaction: *F*_1,36_ = 0.016, *p* = 0.899; Fig. 3d1)(genotype effect: *F*_1,24_ = 0.490, *p* = 0.491; drug effect: *F*_1,24_ = 0.025, *p* = 0.875; genotype X drug interaction: *F*_1,24_ = 0.583, *p* = 0.453; Fig. 3e1). The vehicle-treated *Fmr1* KO mice spent less time in direct interaction with the stranger mouse target (Fig. 3d2 and 3e2). While a single vorinostat administration had no significant effect (genotype effect: *F*_1,36_ = 137.711, *p* = 0.001; drug effect: *F*_1,36_ = 4.386, *p* = 0.043; genotype X drug interaction: *F*_1,36_ = 0.010, *p* = 921; Fig. 3d2), repeated daily vorinostat treatment for 2 weeks improved social interaction in the *Fmr1* KO mice to the WT level (genotype effect: *F*_1,24_ = 29.767, *p* = 0.001; drug effect: *F*_1,24_ = 3.219, *p* = 0.085; genotype X drug interaction: *F*_1,24_ = 8.127, *p* = 0.009; Fig. 3e2).

### Effects of vorinostat on protein synthesis

The abnormally elevated protein synthesis has been recognized as a prominent aspect of cellular pathology associated with FXS ^1,2^. We confirmed that the *Fmr1* KO neurons display a higher level of new protein synthesis than WT neurons (genotype effect: *F*_1,30_ = 13.060, *p* = 0.001; drug effect: *F*_2,30_ = 0.386, *p* = 0.683; genotype X drug interaction: *F*_2,30_ = 0.067, *p* = 0.935; Fig. 4). However, vorinostat failed to suppress protein synthesis in both *Fmr1* KO (*F*_2,30_ = 0.384, *p* = 0.684) and WT neurons (*F*_2,30_ = 0.068, *p* = 0.934) (Fig. 4).

**Figure 4.**
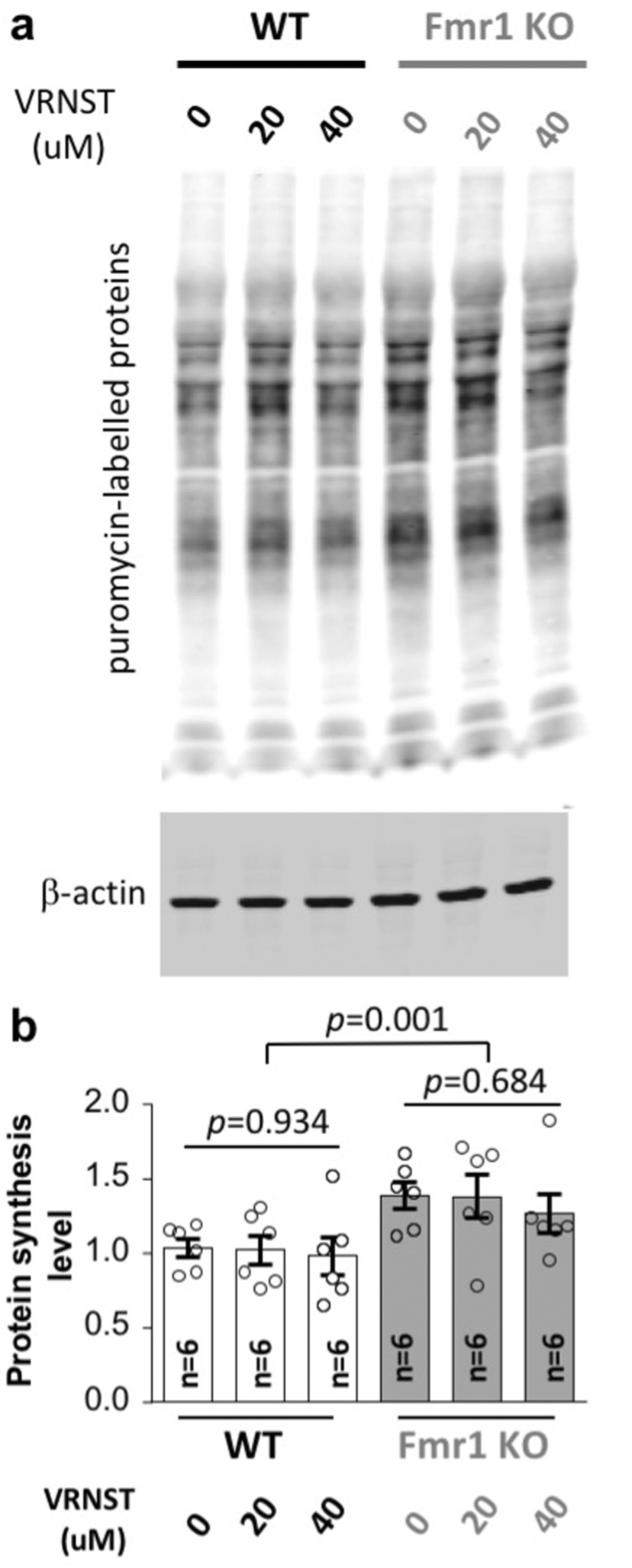
Vorinostat does not correct the elevated protein synthesis in the *Fmr1* KO neurons. Wild type (WT) and *Frm1* KO hippocampal neurons were treated with vehicle or vorinostat (VRNST) at 20 and 40 μM, as indicated, for 30 min, following which puromycin was applied for 30 min. Samples were then collected and subjected to Western blot analysis. Representative images are presented in **a**. Quantification is shown as mean +/- SEM in **b**. The *p*-values were determined by one-way and two-way ANOVA followed by *post hoc* pairwise comparison.

### Trichostatin A does not affect the behavioral outcomes in the *Fmr1* KO mice

As the main known pharmacological action of vorinostat is HDAC inhibition, we wondered whether the observed therapeutic efficacy is due to a general effect of HDAC inhibition and can be achieved by the administration of a different HDAC inhibitor. Trichostatin A is a well recognized HDAC inhibitor ^27^ and a top-ranked similarity drug of vorinostat (Fig. 1). Following vehicle and trichostatin A treatment, we examined WT and *Fmr1* KO mice with object location memory (Fig. 5a) and light-dark box tests (Fig. 5b). Comparing to WT mice, the trichostatin A-treated *Fmr1* KO mice still showed impaired object location memory (location effect: *F*_1,56_ = 0.106, *p* = 0.746; genotype X location interaction: *F*_1,56_ = 0.054, *p* = 0.817; Fig. 5a1) (location effect: *F*_1,56_ = 11.296, *p* = 0.001; genotype X location interaction: *F*_1,56_ = 15.294, *p* = 0.0001; Fig. 5a2) and more transitions between the light and dark chambers (genotype effect: *F*_1,27_ = 24.565, *p* = 0.0001; drug effect: *F*_1,27_ = 1.291, *p* = 0.266; genotype X drug interaction: *F*_1,27_ = 0.025, *p* = 0.875; Fig. 5b1) (genotype effect: *F*_1,27_ = 2.127, *p* = 0.152; drug effect: *F*_1,27_ = 1.437, *p* = 0.241; genotype X drug interaction: *F*_1,27_ = 0.012, *p* = 0.913; Fig. 5b2). We further found that levels of the acetylated histone proteins H2B, H3, and H4 are normal in the *Fmr1* KO neurons (Fig. 5c). These data suggest that the therapeutic effect of vorinostat is unlikely due to its inhibition activity against HDAC.

**Figure 5.**
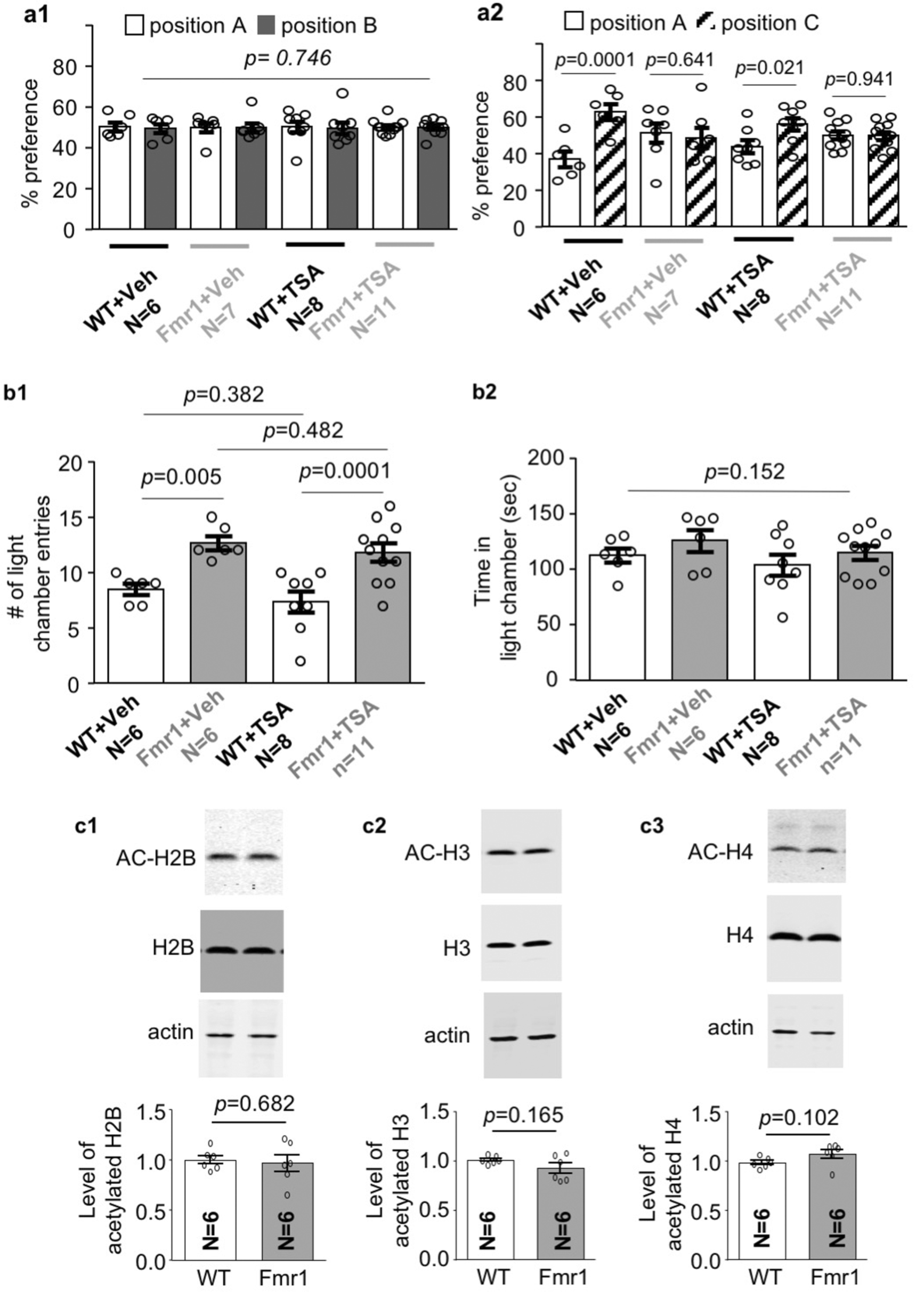
Trichostatin A does not affect object location memory and light/dark box behavior in the *Fmr1* KO mice. a. Wild type (WT) and *Fmr1* KO (Fmr1) mice were injected with vehicle or trichostatin A (TSA). 30 min later, mice were trained to learn object location (**a**) or subjected to a light/dark box test (**b**). In **a**, mouse preference to the objects at locations A and B during training (**a1**) and preference to objects at locations A and C during testing (**a2**) are presented. In **b**, the number of transitions between the light and dark chamber (**b1**) and the time spent in the light chamber (**b2**) are shown. **c**. Samples collected from WT and *Fmr1* KO hippocampal neurons were analyzed for the level of histone acetylation. Western blot was used to determine the level of acetylated H2B, H3, and H4, which were normalized to the total level of the respective histone proteins. Data are presented as mean +/- SEM. The *p*-values were determined three-way (**a**) or two-way (**b**) ANOVA followed by *post hoc* pairwise comparison. Data in **c** were analyzed by Student’s t-test.

## DISCUSSION

While there is no efficacious therapeutics, *de novo* drug development for FXS treatment is still in its infancy and encountered significant obstacles ^28^. One alternative and efficient approach is to repurpose the existing FDA-approved drugs ^29^. In this study, we used the CMap drug-induced transcriptome database to reconfirm vorinostat as a similarity drug to trifluoperazine, which has been recently found to correct FXS-associated symptoms in a mouse model ^12^. The therapeutic efficacy of vorinostat in correcting a variety of behavioral symptoms is validated with the *Fmr1* KO mice.

The use of a holistic analysis of transcriptome signature to predict therapeutic strategy for neurological disorders has been recently proposed but not empirically examined ^25^. One application is to compare the disease-associated transcriptome signature with drug-induced transcriptome signatures. The value of this application is implicated by that transcriptome signature associated with bipolar disorder and schizophrenia can predict drugs, and some of the predicted drugs are already in clinical use to treat these disorders ^24,25^. As the first empirical attempt, we screened the CMap database with the FXS-associated transcriptome signature and identified trifluoperazine as a therapeutics to treat symptoms in the *Fmr1* KO mice ^12^. Another important application of the transcriptome-based therapeutic prediction is to compare the transcriptome signatures associated with different drugs and chemical compounds ^9,10^. Using the trifluoperazine-induced transcriptome change as query identified vorinostat as a top-ranked similarity drug ^12^. In this study, using the vorinostat-induced transcriptome change as a query also identified trifluoperazine as a top-ranked similarity drug. The transcriptome similarity predicts that vorinostat may have similar therapeutic effects to that of trifluoperazine and be useful to treat specific symptoms associated with FXS. Empirically, this study validates the transcriptome-based approach to identify new drug action and repurpose vorinostat.

It is important to note that the vorinostat- and trifluoperazine-induced transcriptome changes share similarities but are not identical. Depending on the cell types, scores (i.e., similarity mean) underlying the similarity between vorinostat and trifluoperazine are 0.282, 0.307, and 0.19 (Fig. 1 and Supplementary Table 1, 2, and 3). As a score of 1 reflects being identical, and a score of -1 reflects being oppositional, it is anticipated that vorinostat and trifluoperazine should have both common and different pharmacological actions. As far as the correction of behavior symptoms is concerned, vorinostat and trifluoperazine have similar but not identical effects. For example, while a single administration of trifluoperazine rescues passive avoidance memory and social deficits ^12^, correction of these deficits requires repeated dosing of vorinostat.

It has been recognized that HDAC inhibitors may be considered to improve learning and memory in animal models of cognition impairment ^15,16,21^. The effect of HDAC inhibitors on non-cognitive functions such as repetitive behavior, hyperactivity, and social interaction has not been recognized and appreciated. Here, we found the therapeutic effects of vorinostat to correct both cognitive and non-cognitive symptoms in *Fmr1* KO mice. Interestingly, vorinostat only affected behavior in the *Fmr1* KO but not WT mice. As the acetylation levels of H2B, H3, and H4 are normal in *Fmr1* KO neurons, it is not straightforward to attribute the therapeutic effect of vorinostat to its HDAC inhibition activity. Further, another HDAC inhibitor trichostatin A failed to rescue deficits of either cognitive or non-cognitive functions. However, we cannot exclude that other HDAC targets are hypo-acetylated in *Fmr1* KO neurons.

Notably, while vorinostat corrects certain FXS-associated behavior symptoms, it does not normalize the elevated protein synthesis in *Fmr1* KO neurons. This is intriguing and suggests that, at least to a certain degree, elevated protein synthesis may not be absolutely linked to all behavioral abnormalities. A recent study found that human FXS samples show various levels of protein synthesis, and fibroblasts derived from some FXS patients display normal translation ^30^. Another complication is that it is not clear whether the elevated translation is universal or restricted to specific brain regions and cell types. Alternatively, vorinostat may dampen the translation of particular FMRP target mRNAs rather than affecting overall protein synthesis. These possibilities remain to be addressed with future studies.

FXS is a complex disorder. It is unlikely that a single treatment strategy will correct all aspects of symptoms. Regarding drug repurposing, several FDA-approved drugs, including minocycline ^31^, metformin ^32^, lovastatin ^33^, and trifluoperazine ^12^ have shown certain therapeutic efficacy in the *Fmr1* KO mice. However, these drugs are not able to rescue all pathological outcomes. Repurposing new therapeutic reagents such as vorinostat will not only provide a new potential treatment choice but also expand the possibility of combination therapy. Regarding the transcriptome-based approach to identify new drug effects, other FDA-approved similarity drugs of trifluoperazine ^12^ and vorinostat (Fig. 1 and Supplementary Tables 1, 2, and 3) may be considered and examined in future studies.

In summary, we used unbiased transcriptome analysis to identify the new therapeutic potential of vorinostat as FXS treatment. We provide evidence to support the value of holistic transcriptome signature in drug repurposing. The effects of vorinostat on correcting cognitive and non-cognitive symptoms in FXS mice encourage human trials.

## AUTHOR CONTRIBUTION

HW initiated the study. XW, XL, and HW identified drug similarity. QD performed empirical investigations and analyzed the data. QD, XL, and HW wrote the manuscript.

## Legends for Supplementary Tables

**Supplementary Table 1** vorinostat similarity compounds screened by DEGs induced by vorinostat in MCF7 cells. The *p*-value determines the ranking of the similarity compounds/drugs; compounds/drugs are listed with their *p*-values in ascending order.

**Supplementary Table 2** vorinostat similarity compounds screened by DEGs induced by vorinostat in PC3 cells. The *p*-value determines the ranking of the similarity compounds/drugs; compounds/drugs are listed with their *p*-values in ascending order.

**Supplementary Table 3** vorinostat similarity compounds screened by DEGs induced by vorinostat in HL60 cells. The *p*-value determines the ranking of the similarity compounds/drugs; compounds/drugs are listed with their *p*-values in ascending order.

## REFERENCES

1. Santoro, M.R., Bray, S.M. & Warren, S.T. Molecular mechanisms of fragile X syndrome: a twenty-year perspective. Annu Rev Pathol 7, 219–245 (2012).

2. Sethna, F., Moon, C. & Wang, H. From FMRP function to potential therapies for fragile X syndrome. Neurochem Res 39, 1016–1031 (2014).

3. Sethna, F., et al. Enhanced expression of ADCY1 underlies aberrant neuronal signalling and behaviour in a syndromic autism model. Nature communications 8, 14359 (2017).

4. Dolen, G., Carpenter, R.L., Ocain, T.D. & Bear, M.F. Mechanism-based approaches to treating fragile X. Pharmacology & therapeutics 127, 78–93 (2010).

5. Gross, C., et al. Isoform-selective phosphoinositide 3-kinase inhibition ameliorates a broad range of fragile X syndrome-associated deficits in a mouse model. Neuropsychopharmacology 44, 324–333 (2019).

6. Bhattacharya, A., et al. Targeting Translation Control with p70 S6 Kinase 1 Inhibitors to Reverse Phenotypes in Fragile X Syndrome Mice. Neuropsychopharmacology 41, 1991–2000 (2016).

7. Toledo, M.A., Wen, T.H., Binder, D.K., Ethell, I.M. & Razak, K.A. Reversal of ultrasonic vocalization deficits in a mouse model of Fragile X Syndrome with minocycline treatment or genetic reduction of MMP-9. Behav Brain Res 372, 112068 (2019).

8. Ashburn, T.T. & Thor, K.B. Drug repositioning: identifying and developing new uses for existing drugs. Nature reviews. Drug discovery 3, 673–683 (2004).

9. Lamb, J., et al. The Connectivity Map: using gene-expression signatures to connect small molecules, genes, and disease. Science 313, 1929–1935 (2006).

10. Qu, X.A. & Rajpal, D.K. Applications of Connectivity Map in drug discovery and development. Drug Discov Today 17, 1289–1298 (2012).

11. Iskar, M., et al. Characterization of drug-induced transcriptional modules: towards drug repositioning and functional understanding. Molecular systems biology 9, 662 (2013).

12. Ding, Q., et al. Transcriptome signature analysis repurposes trifluoperazine for the treatment of fragile X syndrome in mouse model. Communications biology 3, 127 (2020).

13. Falkenberg, K.J. & Johnstone, R.W. Histone deacetylases and their inhibitors in cancer, neurological diseases and immune disorders. Nature reviews. Drug discovery 13, 673–691 (2014).

14. Hockly, E., et al. Suberoylanilide hydroxamic acid, a histone deacetylase inhibitor, ameliorates motor deficits in a mouse model of Huntington’s disease. Proceedings of the National Academy of Sciences of the United States of America 100, 2041–2046 (2003).

15. Benito, E., et al. HDAC inhibitor-dependent transcriptome and memory reinstatement in cognitive decline models. J Clin Invest 125, 3572–3584 (2015).

16. Guan, J.S., et al. HDAC2 negatively regulates memory formation and synaptic plasticity. Nature 459, 55–60 (2009).

17. Irizarry, R.A., et al. Exploration, normalization, and summaries of high density oligonucleotide array probe level data. Biostatistics 4, 249–264 (2003).

18. Gautier, L., Cope, L., Bolstad, B.M. & Irizarry, R.A. affy--analysis of Affymetrix GeneChip data at the probe level. Bioinformatics 20, 307–315 (2004).

19. Ding, Q., Sethna, F. & Wang, H. Behavioral analysis of male and female Fmr1 knockout mice on C57BL/6 background. Behav Brain Res 271, 72–78 (2014).

20. Basu, T., et al. Histone deacetylase inhibitors restore normal hippocampal synaptic plasticity and seizure threshold in a mouse model of Tuberous Sclerosis Complex. Scientific reports 9, 5266 (2019).

21. Sharma, S., Taliyan, R. & Ramagiri, S. Histone deacetylase inhibitor, trichostatin A, improves learning and memory in high-fat diet-induced cognitive deficits in mice. Journal of molecular neuroscience : MN 56, 1–11 (2015).

22. Hsing, C.H., et al. Histone Deacetylase Inhibitor Trichostatin A Ameliorated Endotoxin-Induced Neuroinflammation and Cognitive Dysfunction. Mediators of inflammation 2015, 163140 (2015).

23. Schmidt, E.K., Clavarino, G., Ceppi, M. & Pierre, P. SUnSET, a nonradioactive method to monitor protein synthesis. Nat Methods 6, 275–277 (2009).

24. Gandal, M.J., et al. Shared molecular neuropathology across major psychiatric disorders parallels polygenic overlap. Science 359, 693–697 (2018).

25. So, H.C., et al. Analysis of genome-wide association data highlights candidates for drug repositioning in psychiatry. Nat Neurosci 20, 1342–1349 (2017).

26. Tasic, B., et al. Adult mouse cortical cell taxonomy revealed by single cell transcriptomics. Nat Neurosci 19, 335–346 (2016).

27. Bieliauskas, A.V. & Pflum, M.K. Isoform-selective histone deacetylase inhibitors. Chemical Society reviews 37, 1402–1413 (2008).

28. Berry-Kravis, E.M., et al. Drug development for neurodevelopmental disorders: lessons learned from fragile X syndrome. Nature reviews. Drug discovery 17, 280–299 (2018).

29. Tranfaglia, M.R., et al. Repurposing available drugs for neurodevelopmental disorders: The fragile X experience. Neuropharmacology 147, 74–86 (2019).

30. Jacquemont, S., et al. Protein synthesis levels are increased in a subset of individuals with fragile X syndrome. Human molecular genetics 27, 2039–2051 (2018).

31. Siller, S.S. & Broadie, K. Matrix metalloproteinases and minocycline: therapeutic avenues for fragile X syndrome. Neural plasticity 2012, 124548 (2012).

32. Dy, A.B.C., et al. Metformin as targeted treatment in fragile X syndrome. Clinical genetics 93, 216–222 (2018).

33. Osterweil, E.K., et al. Lovastatin corrects excess protein synthesis and prevents epileptogenesis in a mouse model of fragile X syndrome. Neuron 77, 243–250 (2013).

