## Supplementary Table 1 for "Vorinostat corrects cognitive and non-cognitive symptoms in a mouse model of fragile X syndrome"

| rank | cmap name | cell line | mean | n | enrichment | p-value | percent non-null |
|---|---|---|---|---|---|---|---|
| 1 | trichostatin A | MCF7 | 0.861 | 92 | 0.944 | 0 | 95 |
| 2 | resveratrol | MCF7 | 0.364 | 6 | 0.937 | 0 | 100 |
| 3 | 15-delta prostaglandin J2 | MCF7 | 0.298 | 8 | 0.805 | 0 | 100 |
| 4 | trifluoperazine | MCF7 | 0.282 | 9 | 0.781 | 0 | 88 |
| 5 | thioridazine | MCF7 | 0.259 | 11 | 0.781 | 0 | 90 |
| 6 | tanespimycin | MCF7 | 0.239 | 36 | 0.707 | 0 | 86 |
| 7 | fulvestrant | MCF7 | 0.228 | 21 | 0.683 | 0 | 85 |
| 8 | LY-294002 | MCF7 | 0.234 | 34 | 0.608 | 0 | 73 |
| 9 | valproic acid | MCF7 | 0.233 | 31 | 0.475 | 0 | 54 |
| 10 | moroxydine | MCF7 | -0.826 | 2 | -0.996 | 0.00004 | 100 |
| 11 | lycorine | MCF7 | -0.74 | 3 | -0.978 | 0.00004 | 100 |
| 12 | fluphenazine | MCF7 | 0.163 | 10 | 0.683 | 0.00004 | 70 |
| 13 | troglitazone | MCF7 | 0.256 | 7 | 0.751 | 0.00018 | 85 |
| 14 | phenoxybenzamine | MCF7 | 0.317 | 3 | 0.943 | 0.00022 | 100 |
| 15 | pheneticillin | MCF7 | -0.686 | 2 | -0.989 | 0.00028 | 100 |
| 16 | alvespimycin | MCF7 | 0.243 | 7 | 0.729 | 0.0003 | 85 |
| 17 | racecadotril | MCF7 | -0.693 | 2 | -0.988 | 0.00036 | 100 |
| 18 | prochlorperazine | MCF7 | 0.204 | 9 | 0.64 | 0.00044 | 66 |
| 19 | geldanamycin | MCF7 | 0.198 | 10 | 0.599 | 0.00052 | 70 |
| 20 | demecarium bromide | MCF7 | -0.699 | 2 | -0.983 | 0.00062 | 100 |
| 21 | rottlerin | MCF7 | 0.278 | 3 | 0.922 | 0.00104 | 100 |
| 22 | monastrol | MCF7 | 0.125 | 7 | 0.673 | 0.00104 | 85 |
| 23 | rifabutin | MCF7 | 0.656 | 2 | 0.974 | 0.00107 | 100 |
| 24 | disopyramide | MCF7 | -0.682 | 2 | -0.976 | 0.00125 | 100 |
| 25 | iopromide | MCF7 | -0.696 | 2 | -0.975 | 0.00135 | 100 |
| 26 | dimethadione | MCF7 | -0.646 | 2 | -0.971 | 0.00167 | 100 |
| 27 | syrosingopine | MCF7 | 0.386 | 2 | 0.965 | 0.00211 | 100 |
| 28 | aminophylline | MCF7 | -0.585 | 2 | -0.966 | 0.00254 | 100 |
| 29 | ketorolac | MCF7 | -0.578 | 2 | -0.966 | 0.00254 | 100 |
| 30 | ondansetron | MCF7 | -0.628 | 2 | -0.964 | 0.00292 | 100 |
| 31 | estropipate | MCF7 | -0.693 | 2 | -0.963 | 0.00308 | 100 |
| 32 | arachidonyltrifluoromethane | MCF7 | -0.573 | 2 | -0.959 | 0.00366 | 100 |
| 33 | mycophenolic acid | MCF7 | 0.351 | 2 | 0.951 | 0.00439 | 100 |
| 34 | rescinnamine | MCF7 | 0.326 | 2 | 0.95 | 0.00449 | 100 |
| 35 | thiostrepton | MCF7 | 0.332 | 2 | 0.948 | 0.00497 | 100 |
| 36 | mafenide | MCF7 | -0.583 | 2 | -0.95 | 0.00551 | 100 |
| 37 | nortriptyline | MCF7 | 0.312 | 2 | 0.945 | 0.00567 | 100 |
| 38 | depudecin | MCF7 | 0.344 | 2 | 0.945 | 0.00573 | 100 |
| 39 | bufexamac | MCF7 | 0.35 | 2 | 0.943 | 0.00606 | 100 |
| 40 | raloxifene | MCF7 | 0.277 | 3 | 0.85 | 0.00641 | 100 |
| 41 | butyl hydroxybenzoate | MCF7 | -0.347 | 3 | -0.849 | 0.00681 | 66 |
| 42 | methylbenzethonium chloride | MCF7 | 0.223 | 3 | 0.844 | 0.00749 | 100 |
| 43 | sulconazole | MCF7 | 0.287 | 2 | 0.931 | 0.00891 | 100 |
| 44 | hycanthone | MCF7 | 0.315 | 2 | 0.929 | 0.00972 | 100 |
| 45 | homochlorcyclizine | MCF7 | 0.296 | 2 | 0.927 | 0.01022 | 100 |
| 46 | nicergoline | MCF7 | 0.289 | 2 | 0.927 | 0.01022 | 100 |
| 47 | 5182598 | MCF7 | -0.5 | 2 | -0.929 | 0.0105 | 50 |
| 48 | lobeline | MCF7 | -0.419 | 2 | -0.927 | 0.01095 | 50 |
| 49 | benzethonium chloride | MCF7 | 0.282 | 2 | 0.925 | 0.01097 | 100 |
| 50 | mesoridazine | MCF7 | -0.333 | 2 | -0.927 | 0.01099 | 50 |
| 51 | pimethixene | MCF7 | 0.279 | 2 | 0.923 | 0.01171 | 100 |

| | | | | | | | |
|---|---|---|---|---|---|---|---|
| 52 | monorden | MCF7 | 0.156 | 12 | 0.44 | 0.01203 | 58 |
| 53 | carcinine | MCF7 | -0.39 | 2 | -0.923 | 0.01205 | 50 |
| 54 | clomifene | MCF7 | 0.295 | 2 | 0.922 | 0.01217 | 100 |
| 55 | thioproperazine | MCF7 | 0.271 | 2 | 0.921 | 0.01223 | 100 |
| 56 | quinostatin | MCF7 | 0.279 | 2 | 0.918 | 0.01314 | 100 |
| 57 | 5707885 | MCF7 | 0.27 | 2 | 0.918 | 0.01336 | 100 |
| 58 | fluspirilene | MCF7 | 0.267 | 2 | 0.917 | 0.01364 | 100 |
| 59 | cefixime | MCF7 | -0.445 | 2 | -0.915 | 0.01453 | 50 |
| 60 | protriptyline | MCF7 | 0.289 | 2 | 0.913 | 0.01511 | 100 |
| 61 | harmol | MCF7 | -0.319 | 2 | -0.913 | 0.01529 | 50 |
| 62 | ciclopirox | MCF7 | 0.266 | 2 | 0.912 | 0.01541 | 100 |
| 63 | CP-645525-01 | MCF7 | 0.267 | 2 | 0.912 | 0.01559 | 100 |
| 64 | cyclopenthiazide | MCF7 | -0.347 | 2 | -0.911 | 0.01589 | 50 |
| 65 | desipramine | MCF7 | 0.289 | 2 | 0.91 | 0.0163 | 100 |
| 66 | clomipramine | MCF7 | 0.265 | 2 | 0.91 | 0.01648 | 100 |
| 67 | mefloquine | MCF7 | 0.259 | 2 | 0.91 | 0.01664 | 100 |
| 68 | suprofen | MCF7 | -0.298 | 2 | -0.909 | 0.0167 | 50 |
| 69 | zidovudine | MCF7 | -0.326 | 2 | -0.908 | 0.01686 | 50 |
| 70 | apramycin | MCF7 | -0.33 | 2 | -0.906 | 0.01787 | 50 |
| 71 | prenylamine | MCF7 | 0.25 | 2 | 0.906 | 0.01813 | 100 |
| 72 | amoxapine | MCF7 | 0.282 | 2 | 0.905 | 0.01837 | 100 |
| 73 | withaferin A | MCF7 | 0.257 | 2 | 0.905 | 0.01853 | 100 |
| 74 | epivincamine | MCF7 | -0.395 | 2 | -0.903 | 0.01865 | 50 |
| 75 | perhexiline | MCF7 | 0.259 | 2 | 0.903 | 0.0194 | 100 |
| 76 | (+/-)-catechin | MCF7 | -0.281 | 2 | -0.901 | 0.01966 | 50 |
| 77 | nitrofurantoin | MCF7 | 0.25 | 2 | 0.901 | 0.02026 | 100 |
| 78 | cinoxacin | MCF7 | -0.328 | 2 | -0.898 | 0.02072 | 50 |
| 79 | loperamide | MCF7 | 0.179 | 3 | 0.783 | 0.02097 | 66 |
| 80 | parthenolide | MCF7 | 0.253 | 2 | 0.898 | 0.02167 | 100 |
| 81 | dosulepin | MCF7 | 0.247 | 2 | 0.897 | 0.02195 | 100 |
| 82 | colforsin | MCF7 | 0.271 | 2 | 0.895 | 0.02272 | 100 |
| 83 | talampicillin | MCF7 | -0.336 | 2 | -0.894 | 0.02276 | 50 |
| 84 | nordihydroguaiaretic acid | MCF7 | 0.164 | 8 | 0.499 | 0.02319 | 62 |
| 85 | 0317956-0000 | MCF7 | 0.103 | 4 | 0.674 | 0.02485 | 50 |
| 86 | corticosterone | MCF7 | 0.263 | 2 | 0.889 | 0.02559 | 100 |
| 87 | terfenadine | MCF7 | 0.255 | 2 | 0.887 | 0.0263 | 100 |
| 88 | etoposide | MCF7 | 0.29 | 2 | 0.886 | 0.02702 | 100 |
| 89 | 5224221 | MCF7 | 0.252 | 2 | 0.885 | 0.0273 | 100 |
| 90 | tonzonium bromide | MCF7 | 0.229 | 2 | 0.883 | 0.02811 | 100 |
| 91 | pentoxyverine | MCF7 | -0.236 | 2 | -0.881 | 0.02831 | 50 |
| 92 | meclocycline | MCF7 | -0.452 | 2 | -0.879 | 0.02931 | 50 |
| 93 | metanephrine | MCF7 | 0.22 | 2 | 0.879 | 0.02936 | 100 |
| 94 | esculin | MCF7 | -0.246 | 2 | -0.87 | 0.03356 | 50 |
| 95 | pizotifen | MCF7 | 0.217 | 2 | 0.869 | 0.03479 | 100 |
| 96 | bepridil | MCF7 | 0.214 | 2 | 0.867 | 0.03608 | 100 |
| 97 | alimemazine | MCF7 | 0.219 | 2 | 0.867 | 0.0364 | 100 |
| 98 | NS-398 | MCF7 | 0.231 | 2 | 0.866 | 0.03672 | 100 |
| 99 | butoconazole | MCF7 | 0.228 | 2 | 0.865 | 0.0369 | 100 |
| 100 | cefuroxime | MCF7 | -0.38 | 2 | -0.864 | 0.03696 | 50 |
| 101 | pyrvinium | MCF7 | 0.183 | 4 | 0.647 | 0.03724 | 75 |
| 102 | digoxigenin | MCF7 | 0.122 | 3 | 0.733 | 0.03754 | 66 |
| 103 | clenbuterol | MCF7 | -0.36 | 3 | -0.736 | 0.03768 | 66 |

| | | | | | | | |
|---|---|---|---|---|---|---|---|
| 104 | CP-690334-01 | MCF7 | 0.258 | 4 | 0.644 | 0.03867 | 75 |
| 105 | probucol | MCF7 | -0.277 | 4 | -0.637 | 0.04201 | 50 |
| 106 | disulfiram | MCF7 | 0.224 | 2 | 0.855 | 0.04284 | 100 |
| 107 | cinchocaine | MCF7 | 0.194 | 2 | 0.846 | 0.0476 | 100 |
| 108 | sulfathiazole | MCF7 | -0.374 | 2 | -0.842 | 0.04996 | 50 |
