## Supplementary Table 2 for "Vorinostat corrects cognitive and non-cognitive symptoms in a mouse model of fragile X syndrome"

| rank | cmap name | cell line | mean | n | enrichment | p-value | percent non-null |
|---|---|---|---|---|---|---|---|
| 1 | trichostatin A | PC3 | 0.81 | 55 | 0.976 | 0 | 98 |
| 2 | thioridazine | PC3 | 0.289 | 5 | 0.925 | 0.00002 | 100 |
| 3 | LY-294002 | PC3 | 0.181 | 12 | 0.665 | 0.00002 | 75 |
| 4 | Prestwick-675 | PC3 | -0.909 | 2 | -0.998 | 0.00004 | 100 |
| 5 | scriptaid | PC3 | 0.772 | 2 | 0.992 | 0.00006 | 100 |
| 6 | AH-6809 | PC3 | -0.756 | 2 | -0.995 | 0.00008 | 100 |
| 7 | trifluoperazine | PC3 | 0.307 | 3 | 0.96 | 0.0001 | 100 |
| 8 | lidocaine | PC3 | -0.681 | 2 | -0.99 | 0.00026 | 100 |
| 9 | fluphenazine | PC3 | 0.27 | 3 | 0.942 | 0.00026 | 100 |
| 10 | valproic acid | PC3 | 0.257 | 10 | 0.595 | 0.0006 | 70 |
| 11 | calcium folinate | PC3 | -0.612 | 2 | -0.978 | 0.00101 | 100 |
| 12 | MS-275 | PC3 | 0.545 | 2 | 0.975 | 0.00101 | 100 |
| 13 | 15-delta prostaglandin J2 | PC3 | 0.271 | 3 | 0.92 | 0.00112 | 100 |
| 14 | MG-262 | PC3 | 0.343 | 2 | 0.968 | 0.00173 | 100 |
| 15 | josamycin | PC3 | -0.574 | 2 | -0.971 | 0.00175 | 100 |
| 16 | astemizole | PC3 | 0.333 | 2 | 0.965 | 0.00207 | 100 |
| 17 | beta-escin | PC3 | 0.319 | 2 | 0.964 | 0.00221 | 100 |
| 18 | mefloquine | PC3 | 0.296 | 2 | 0.962 | 0.00256 | 100 |
| 19 | dilazep | PC3 | 0.332 | 2 | 0.96 | 0.00266 | 100 |
| 20 | helveticoside | PC3 | 0.307 | 2 | 0.959 | 0.00294 | 100 |
| 21 | withaferin A | PC3 | 0.278 | 2 | 0.957 | 0.0033 | 100 |
| 22 | azacyclonol | PC3 | 0.271 | 2 | 0.955 | 0.00366 | 100 |
| 23 | methylbenzethonium chloride | PC3 | 0.284 | 2 | 0.953 | 0.00404 | 100 |
| 24 | resveratrol | PC3 | 0.271 | 2 | 0.952 | 0.00416 | 100 |
| 25 | perphenazine | PC3 | 0.262 | 2 | 0.948 | 0.00497 | 100 |
| 26 | prochlorperazine | PC3 | 0.254 | 3 | 0.859 | 0.00531 | 100 |
| 27 | mianserin | PC3 | 0.249 | 2 | 0.942 | 0.00638 | 100 |
| 28 | disulfiram | PC3 | 0.262 | 2 | 0.941 | 0.00672 | 100 |
| 29 | cloperastine | PC3 | 0.232 | 2 | 0.935 | 0.00817 | 100 |
| 30 | alexidine | PC3 | 0.248 | 2 | 0.934 | 0.00819 | 100 |
| 31 | lomustine | PC3 | 0.233 | 2 | 0.929 | 0.00962 | 100 |
| 32 | loperamide | PC3 | 0.219 | 2 | 0.92 | 0.01272 | 100 |
| 33 | captopril | PC3 | -0.337 | 2 | -0.916 | 0.01433 | 50 |
| 34 | CP-690334-01 | PC3 | 0.2 | 4 | 0.708 | 0.01494 | 50 |
| 35 | PHA-00846566E | PC3 | -0.415 | 2 | -0.912 | 0.01553 | 50 |
| 36 | famotidine | PC3 | -0.322 | 2 | -0.91 | 0.01638 | 50 |
| 37 | spironolactone | PC3 | 0.211 | 2 | 0.91 | 0.01642 | 100 |
| 38 | nomifensine | PC3 | -0.334 | 2 | -0.909 | 0.01674 | 50 |
| 39 | bromopride | PC3 | -0.315 | 2 | -0.904 | 0.01855 | 50 |
| 40 | chlorcyclizine | PC3 | 0.209 | 2 | 0.903 | 0.0194 | 100 |
| 41 | 5194442 | PC3 | 0.201 | 2 | 0.899 | 0.0207 | 100 |
| 42 | AR-A014418 | PC3 | -0.397 | 2 | -0.898 | 0.02072 | 50 |
| 43 | riluzole | PC3 | -0.389 | 2 | -0.896 | 0.02157 | 50 |
| 44 | hydralazine | PC3 | -0.294 | 2 | -0.894 | 0.02235 | 50 |
| 45 | SC-19220 | PC3 | -0.436 | 2 | -0.893 | 0.02304 | 50 |
| 46 | tolfenamic acid | PC3 | -0.327 | 2 | -0.882 | 0.02803 | 50 |
| 47 | PHA-00816795 | PC3 | -0.45 | 2 | -0.882 | 0.02819 | 50 |
| 48 | mebendazole | PC3 | 0.206 | 2 | 0.882 | 0.02831 | 100 |
| 49 | 0198306-0000 | PC3 | -0.402 | 2 | -0.879 | 0.02925 | 50 |
| 50 | antimycin A | PC3 | 0.217 | 2 | 0.877 | 0.03026 | 100 |
| 51 | CP-863187 | PC3 | -0.396 | 2 | -0.875 | 0.03109 | 50 |

| | | | | | | | |
|---|---|---|---|---|---|---|---|
| 52 | hydrastinine | PC3 | -0.321 | 2 | -0.871 | 0.03278 | 50 |
| 53 | PNU-0293363 | PC3 | -0.303 | 2 | -0.866 | 0.03626 | 50 |
| 54 | 16,16-dimethylprostaglandin E2 | PC3 | 0.188 | 2 | 0.866 | 0.03648 | 100 |
| 55 | troglitazone | PC3 | 0.153 | 4 | 0.647 | 0.03712 | 75 |
| 56 | acenocoumarol | PC3 | -0.372 | 2 | -0.86 | 0.03899 | 50 |
| 57 | orphenadrine | PC3 | 0.215 | 2 | 0.861 | 0.03901 | 100 |
| 58 | ethosuximide | PC3 | -0.372 | 2 | -0.855 | 0.04191 | 50 |
| 59 | oxetacaine | PC3 | 0.188 | 2 | 0.856 | 0.04231 | 100 |
| 60 | gallamine triethiodide | PC3 | -0.293 | 2 | -0.854 | 0.04241 | 50 |
| 61 | tropicamide | PC3 | -0.315 | 2 | -0.852 | 0.0437 | 50 |
| 62 | triamcinolone | PC3 | -0.35 | 2 | -0.845 | 0.04825 | 50 |
