## Supplementary Table 3 for "Vorinostat corrects cognitive and non-cognitive symptoms in a mouse model of fragile X syndrome"

| rank | cmap name | cell line | mean | n | enrichment | p-value | percent non-null |
|---|---|---|---|---|---|---|---|
| 1 | trichostatin A | HL60 | 0.573 | 34 | 0.965 | 0 | 100 |
| 2 | valproic acid | HL60 | 0.423 | 14 | 0.81 | 0 | 85 |
| 3 | LY-294002 | HL60 | 0.291 | 13 | 0.793 | 0 | 84 |
| 4 | tanespimycin | HL60 | 0.187 | 12 | 0.759 | 0 | 83 |
| 5 | 15-delta prostaglandin J2 | HL60 | 0.351 | 3 | 0.964 | 0.00006 | 100 |
| 6 | genistein | HL60 | 0.297 | 3 | 0.96 | 0.0001 | 100 |
| 7 | geldanamycin | HL60 | 0.242 | 3 | 0.942 | 0.00024 | 100 |
| 8 | tretinoin | HL60 | 0.184 | 5 | 0.806 | 0.00062 | 80 |
| 9 | fluphenazine | HL60 | 0.102 | 4 | 0.749 | 0.0076 | 50 |
| 10 | trifluoperazine | HL60 | 0.19 | 4 | 0.742 | 0.00855 | 75 |
| 11 | sirolimus | HL60 | 0.102 | 10 | 0.494 | 0.00907 | 50 |
| 12 | prochlorperazine | HL60 | 0.192 | 4 | 0.724 | 0.0116 | 75 |
| 13 | thioridazine | HL60 | 0.184 | 4 | 0.695 | 0.0181 | 75 |
| 14 | troglitazone | HL60 | 0.163 | 4 | 0.678 | 0.02335 | 75 |
| 15 | raloxifene | HL60 | 0.158 | 2 | 0.888 | 0.02622 | 100 |
| 16 | sodium phenylbutyrate | HL60 | 0.197 | 2 | 0.886 | 0.02694 | 100 |
